# Dynamic cell cycle-dependent phosphorylation modulates CENP-L-CENP-N centromere recruitment

**DOI:** 10.1101/2021.12.17.473210

**Authors:** Alexandra P. Navarro, Iain M. Cheeseman

## Abstract

The kinetochore is a macromolecular structure that is required to ensure proper chromosome segregation during each cell division. The kinetochore is assembled upon a platform of the 16-subunit Constitutive Centromere Associated Network (CCAN), which is present at centromeres throughout the cell cycle. The nature and regulation of CCAN assembly, interactions, and dynamics required to facilitate changing centromere properties and requirements remain to be fully elucidated. The CENP-LN CCAN sub-complex displays a unique cell cycle-dependent localization behavior, peaking in S phase. Here, we demonstrate that phosphorylation of CENP-L and CENP-N controls CENP-LN complex formation and localization in a cell cycle-dependent manner. Mimicking constitutive phosphorylation of either CENP-L or CENP-N or simultaneously preventing phosphorylation of both proteins prevents CENP-LN localization and disrupts chromosome segregation. Together, our work suggests that cycles of phosphorylation and dephosphorylation are critical for CENP-LN complex recruitment and dynamics at centromeres to enable cell cycle-dependent CCAN reorganization.

## Introduction

The kinetochore plays a critical role in ensuring the fidelity of chromosome segregation during each cell division (Cheeseman, 2014). Kinetochore assembly is restricted to the centromere of each chromosome due to the presence of a centromere-specific histone H3 variant called CENP-A. Within the inner kinetochore, the 16-subunit complex Constitutive Centromere-Associated Network (CCAN) acts to recognize CENP-A. These inner kinetochore proteins are defined by their constitutive assembly at centromeres throughout all phases of the cell cycle (Navarro and Cheeseman, 2021). The CCAN, in turn, provides a platform for assembly of the outer kinetochore, which ultimately binds to spindle microtubules to facilitate chromosome segregation. The proteins of the CCAN are canonically grouped into 5 sub-complexes: CENP-C; CENP-LN; CENP-HIKM; CENP-TWSX; and CENP-OPQUR (Amano et al., 2009; Basilico et al., 2014; Carroll et al., 2009; Earnshaw and Rothfield, 1985; Foltz et al., 2006; Hori et al., 2008b; Nishino et al., 2012; Okada et al., 2006; Saitoh et al., 1992; Weir et al., 2016). Of these proteins, CENP-C and CENP-N recognize and bind to CENP-A directly (Allu et al., 2019; Carroll et al., 2009; Falk et al., 2015; Guse et al., 2011; Kato et al., 2013). CCAN assembly relies on the interdependent interactions between these protein complexes such that the absence of one protein at centromeres results in the mis-localization of other inner kinetochore sub-complexes (Basilico et al., 2014; Hori et al., 2008a; McKinley et al., 2015; Okada et al., 2006).

Although the inner kinetochore is localized constitutively to centromeres, the structure and organization of this protein assembly changes in a cell cycle-dependent manner (Navarro and Cheeseman, 2020). First, centromeres undergo a significant structural change during G2 and upon entry into mitosis when inner kinetochore proteins CENP-C and CENP-T serve as a binding site for outer kinetochore recruitment (Cheeseman et al., 2008; Gascoigne and Cheeseman, 2013; Malvezzi et al., 2013). Outer kinetochore assembly is restricted to mitosis due to the requirement for phosphorylation of CENP-C and CENP-T by mitotic kinases. (Gascoigne et al., 2011; Huis In ‘t Veld et al., 2016; Pekgoz Altunkaya et al., 2016; Screpanti et al., 2011). Coupled with this change in kinetochore architecture, the interdependent relationships between CCAN subcomplexes also change. For example, during interphase CENP-C relies on its interaction with CENP-LN and CENP-HIKM for its centromere localization, whereas during mitosis CENP-C can localize to kinetochores in the absence of other CCAN subcomplexes (McKinley et al., 2015; Nagpal et al., 2015). This change is mediated in part by the phosphorylation of CENP-C in mitosis by cyclin-dependent kinase (CDK), which acts to increase the affinity of CENP-C for CENP-A and thereby promote its association with kinetochores independently of its association with other CCAN components (Watanabe et al., 2019). Together, these regulated changes in kinetochore organization indicate that a molecular reorganization occurs within the inner kinetochore between interphase and mitosis. However, how the other proteins of the inner kinetochore adapt to these changes has not been fully elucidated.

The CENP-L/N complex plays a critical role in the assembly of the inner kinetochore as it binds directly to both CENP-A and other CCAN components. Although CENP-N and CENP-L are thought to interact stably, each protein performs a specific function within this complex. The N-terminus of CENP-N interacts directly with a distinct loop within CENP-A and this interaction is sufficient to recruit CENP-N to centromeres during interphase (Carroll et al., 2010; Carroll et al., 2009; Fang et al., 2015; Guo et al., 2017; McKinley et al., 2015; Pentakota et al., 2017). The C-terminus of CENP-N interacts directly with CENP-L and through this interaction binds to CENP-C and CENP-HIKM (Carroll et al., 2009; Hinshaw and Harrison, 2013; McKinley et al., 2015; Pentakota et al., 2017). Together, CENP-N and CENP-L effectively bridge the interface between centromere DNA and the assembled CCAN. However, CENP-N levels at kinetochores decrease in mitosis. Recent structural models have suggested that mitotic kinetochores contain sub-stoichiometric amounts of the CENP-N relative to interphase (Allu et al., 2019; Fang et al., 2015; Hellwig et al., 2008). Here, we show that CENP-N and CENP-L are phosphorylated in mitosis and that phosphorylation of either protein is sufficient to prevent kinetochore localization and disrupt the interaction between CENP-N and CENP-L. The dynamic nature of this phosphorylation event is critical, as mutants that fully prevent CENP-LN complex phosphorylation also disrupt its kinetochore recruitment. This work highlights the molecular changes that occur within the inner kinetochore throughout the cell cycle and helps reveal the features that enable a functional mitotic kinetochore assembly state.

## Results and Discussion

### CENP-L/N kinetochore localization decreases upon entry into mitosis

To assess changes in the organization of the inner kinetochore throughout the cell cycle, we sought to quantify the localization of inner kinetochore components that interact directly with the centromere histone, CENP-A (Fig. 1A). Based on antibody staining, we found that CENP-C levels remain relatively constant throughout the cell cycle (Fig. 1B, 1C), consistent with prior work (Gascoigne and Cheeseman, 2013). Prior studies analyzing endogenous CENP-N localization have been limited by the nature of the available CENP-N antibodies. Therefore, we assessed CENP-N localization in cells stably expressing N-terminally tagged GFP-CENP-N as the only CENP-N species present using a CRISPR/Cas9-based replacement strategy to deplete endogenous CENP-N (see (McKinley et al., 2015)) or using C-terminally tagged CENP-N tagged at its endogenous locus. In contrast to CENP-C, CENP-N was enriched during S phase, but then decreased as cells progressed to mitosis (Fig. 1B, 1C), consistent with prior work (Allu et al., 2019; Fang et al., 2015; Hellwig et al., 2011). Within the CCAN, CENP-N forms an obligate complex with CENP-L (Fig. 1A). To assess CENP-L levels, we utilized a similar replacement assay with a GFP-CENP-L fusion. Similar to CENP-N, we found that CENP-L was enriched at kinetochores in S phase followed by reduction to 30% relative of its maximal enrichment upon entry into mitosis (Fig. 1B, 1C). The increased levels of CENP-N and CENP-L at S phase kinetochores coincides with the cell cycle stage during which new molecules of CENP-N are recruited to centromeres (Fang et al., 2015; Hellwig et al., 2011). Therefore, whereas CENP-C levels remain constant throughout the cell cycle, CENP-N and CENP-L kinetochore localization is enriched at S phase but reduced upon entry into mitosis.

**Figure 1.**
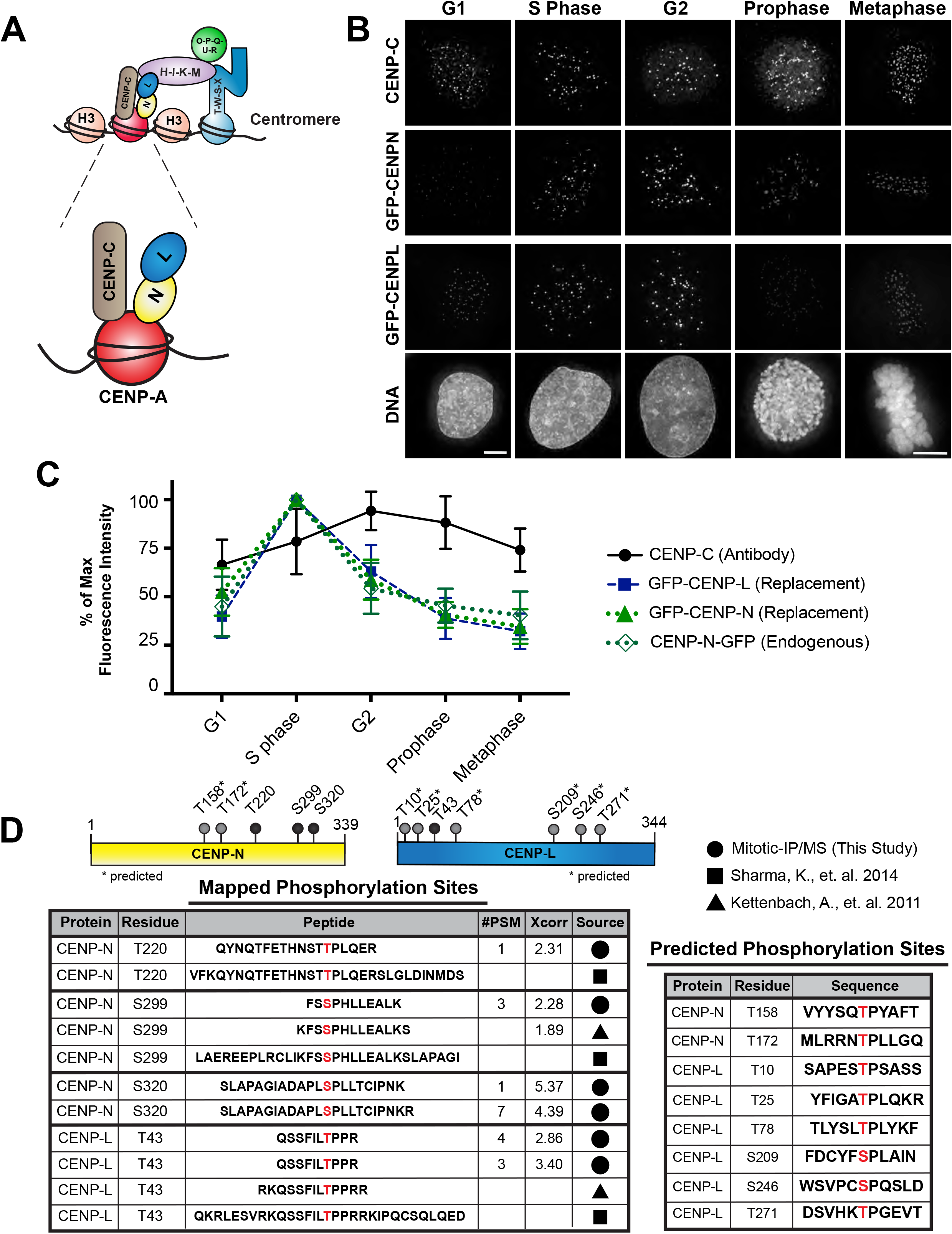
CENP-N and CENP-L levels vary throughout the cell cycle. **A.** Diagram illustrating the proteins that interact directly with centromere histone, CENP-A, within the CCAN. **B.** Representative images of protein levels throughout the cell cycle. All images are scaled the same. DNA images shown in this figure are from the GFP-CENP-L cells shown in this image. Scale bar is 5 μm. **C.** Quantification of CENP-C, CENP-N, and CENP-L levels throughout the cell cycle. CENP-C levels were detected with a CENP-C antibody. Cell cycle stages were determine using the following markers: G1/prophase/mitosis-DM1a; S phase-PCNA; G2-Cyclin B. Levels are presented as a percent of max fluorescence. Quantification of GFP-CENP-N levels throughout the cell cycle. Levels are presented as a percent of max fluorescence. Quantification of GFP-CENP-L levels throughout the cell cycle. Levels are presented as a percent of max fluorescence. **D.** Tables representing the mapped and predicted phosphorylation sites for CENP-N and CENP-L. Phosphorylation sites mapped by mass spec include results from my data and previously published data of mitotically enriched proteome wide mass spec analysis. Predicted phosphorylation sites represent sides that were targeted due to their alignment with the minimal CDK consensus sequence [S/T]-P. All sites were mutated in each construct to generate phosphomutants.

### CENP-L and CENP-N are phosphorylated in mitosis

This altered localization of the CENP-L/N complex in mitosis suggests that this complex undergoes cell cycle specific changes that alter its centromere localization. Based on the prevalence of post-translational modifications in regulating diverse aspects of kinetochore assembly and function (Hara and Fukagawa, 2020; Navarro and Cheeseman, 2020), we hypothesized that CENP-N and CENP-L might be post-translationally modified in a cell cycle-dependent manner. To test this, we isolated GFP-CENP-N from cells enriched in either mitosis or in S phase and GFP-CENP-L from mitotically enriched cells using immuno-precipitation. Our mass spectrometry analysis identified established interacting partners for CENP-N and CENP-L (Supplemental Fig. 1A, 1B). In addition, we identified phosphorylation sites in CENP-N and CENP-L that were present specifically in samples isolated from mitotic cells (Fig. 1D; Supplemental Fig. 1C). These sites have also been identified in proteome-wide mass spectrometry studies of mitotic cells (Hornbeck et al., 2015; Kettenbach et al., 2011; Sharma et al., 2014) (Fig. 1D). We note that the sequence composition of CENP-N and CENP-L precludes the ability to identify peptides spanning their entire sequence using standard mass spectrometry approaches (see PeptideAtlas - https://db.systemsbiology.net/), such that there are likely to be additional phosphorylation events beyond those identified here and in prior work. Interestingly, the majority of the phosphorylation sites identified in CENP-N and CENP-L correspond to a minimal cyclin-dependent kinase (CDK) consensus sequence (pS/pT-[P]; pS/pT-[P]-X-R) (Songyang et al., 1994) (Fig. 1D). Together, our mass spec data indicate that both CENP-N and CENP-L are phosphorylated in mitosis.

### Phosphorylation affects CENP-N and CENP-L function

We next sought to determine the consequences of CENP-N and CENP-L phosphorylation by generating phospho-mutants for both proteins. For these studies, we mutated all CDK consensus phosphorylation sites present in either protein (Fig. 1D, 2A, 2D). Importantly, many of these SP/TP sites are conserved in other vertebrate CENP-L and CENP-N orthologs (Supplemental Fig. 1C, 1D). To test the consequence of preventing phosphorylation, we mutated each serine or threonine residue to alanine (CDK-SA mutant). Reciprocally, to mimic constitutive phosphorylation at these CDK sites, we mutated each serine or threonine to aspartate (CDK-SD mutant). Depletion of CENP-N and CENP-L using a Cas9-inducible knockout strategy with a single guide RNA (sgRNA) targeting the corresponding gene results in severe chromosome misalignment (McKinley et al., 2015) (Fig. 2B, 2C). However, this phenotype was rescued by the expression of wild type GFP-tagged transgenes hardened against the corresponding sgRNA sequence (Fig. 2B-C, 2E-F). Similarly, we found that CDK-SA phospho-null mutants of either CENP-N or CENP-L localized to kinetochores and were able to support proper chromosome alignment and segregation in the absence of the endogenous protein, with only modest mitotic phenotypes (Fig. 2B-C, 2E-F). However, in the presence of the endogenous CENP-N protein, we found that the CENP-N^CDK-SA^ mutant displayed substantially reduced kinetochore localization during both interphase and mitosis (Supplemental Fig. 2A). This suggests that the CENP-N^CDK-SA^ mutant cannot fully compete with the endogenous protein for its localization to the kinetochore. In contrast to these relatively modest phenotypes in the absence of CDK phosphorylation, we found that mutants mimicking the constitutively phosphorylated state (CDK-SD) displayed substantial mitotic defects. Both CENP-N^CDKSD^ and CENP-L^CDKSD^ were unable to localize to kinetochores and failed to rescue proper chromosome segregation (Fig. 2B-C, 2E-F). Similar studies testing phospho-mutants that prevented phosphorylation for the non-CDK consensus phosphorylation sites did not result in changes in localization or function (Supplemental Fig. 1C; data not shown). Together, this data suggests that the phosphorylation of either CENP-L or CENP-N downstream of CDK affects their ability to localize to kinetochores such that mimicking constitutive phosphorylation disrupts kinetochore function for either protein.

**Figure 2.**
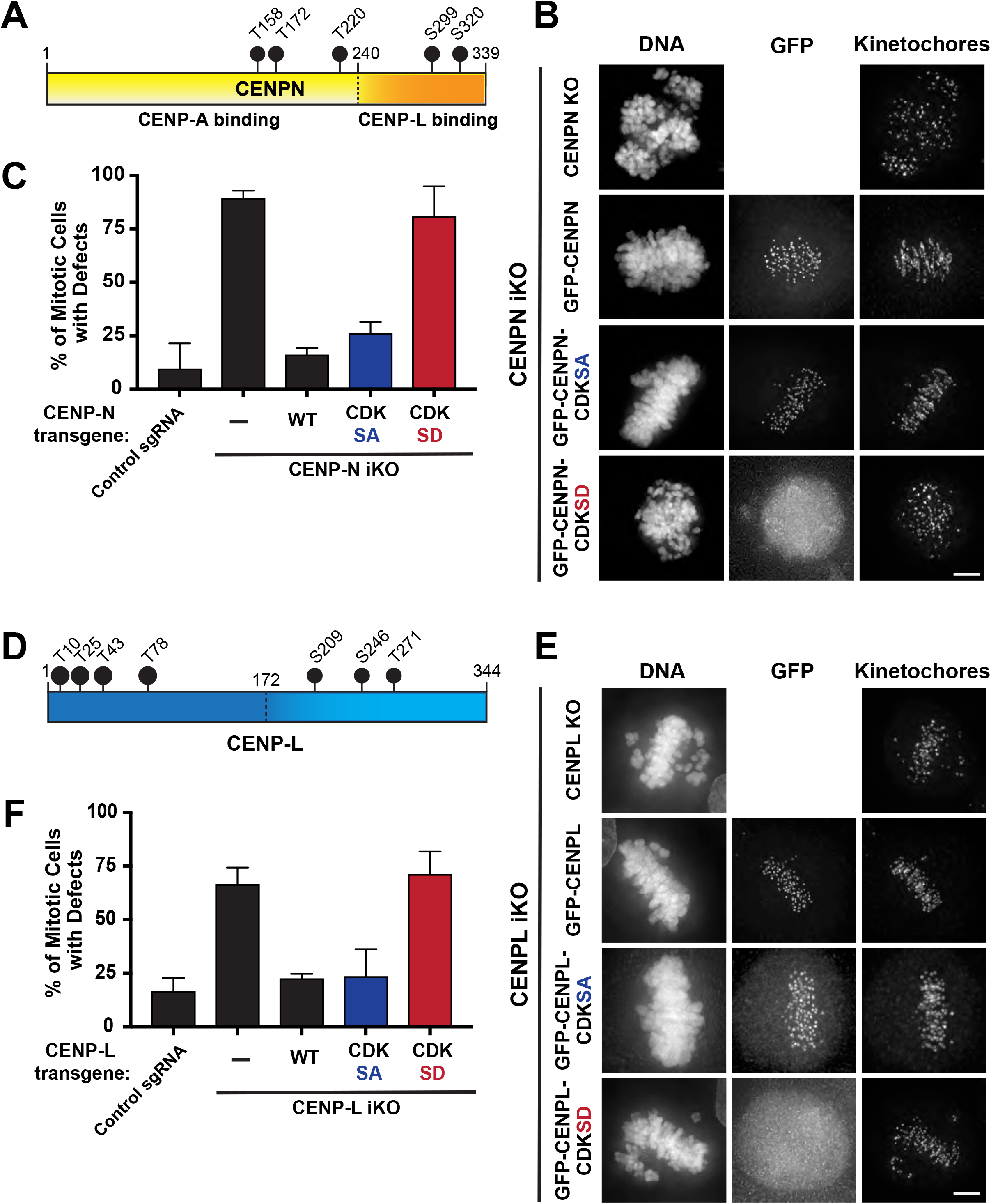
Mutating phosphorylation sites in CENP-L and CENP-N affect their ability to localize to the kinetochore. **A.** Diagram illustrating the SP/TP sites in CENP-N. **B.** Representative images of mitotic phenotypes for each replacement condition. All images were deconvolved. Images are not scaled the same. Scale bar is 5 μm. **C.** Quantification of mitotic phenotypes in CENP-N replacement cell lines after 5 day induction of endogenous CENP-N knockout. Approximately 100 cells were counted for each condition. **D.** Diagram illustrating the SP/TP sites in CENP-L. **E.** Representative images of mitotic phenotypes for each replacement condition. All images were deconvolved. Images are not scaled the same. Scale bar is 5 μm. **F.** Quantification of mitotic phenotypes in CENP-L replacement cell lines after 5 day induction of endogenous CENP-L knockout. Approximately 100 cells were counted for each condition.

### Phosphorylation affects the interaction between CENP-L and CENP-N

Previous work has found that CENP-L and CENP-N are interdependent for their localization to kinetochores (McKinley et al., 2015; Pentakota et al., 2017). Given that phospho-mimetic versions of CENP-L or CENP-N prevented their kinetochore localization (Fig. 2B, 2E), we hypothesized that phosphorylation might alter the formation of the CENP-LN complex by disrupting their ability to interact. To test this, we took a cell-based approach by adapting a previously established *lacO*-tethering assay (Gascoigne et al., 2011). For this assay, we utilized a U20S cell line that has a stably integrated *lacO* array on chromosome 1 (Gascoigne et al., 2011; Janicki et al., 2004). We then transiently co-expressed a bait construct tagged with LacI-GFP that binds with high affinity to the *lacO* array and a TdTomato-tagged test construct that can only localize to the *lacO* array if it binds to the bait protein. Co-localization of GFP and TdTomato to the *lacO* array serves as a proxy for a molecular interaction (Fig. 3A). As expected, we observed robust recruitment of CENP-L when wild type LacI-GFP-CENP-N was targeted to the *lacO* array (Fig. 3B-C). Reciprocally, LacI-GFP-CENP-L recruited CENP-N to the *lacO* array (Fig. 3D). Preventing CENP-L or CENP-N phosphorylation using phospho-null mutants only modestly altered their interactions in this lacO tethering assay (Fig. 3B-3D). In contrast, phospho-mimetic versions of either CENP-N or CENP-L were unable to recruit the reciprocal protein (Fig. 3B-D). We conclude that mimicking phosphorylation of either CENP-N or CENP-L is sufficient to prevent the interaction between these two proteins.

**Figure 3.**
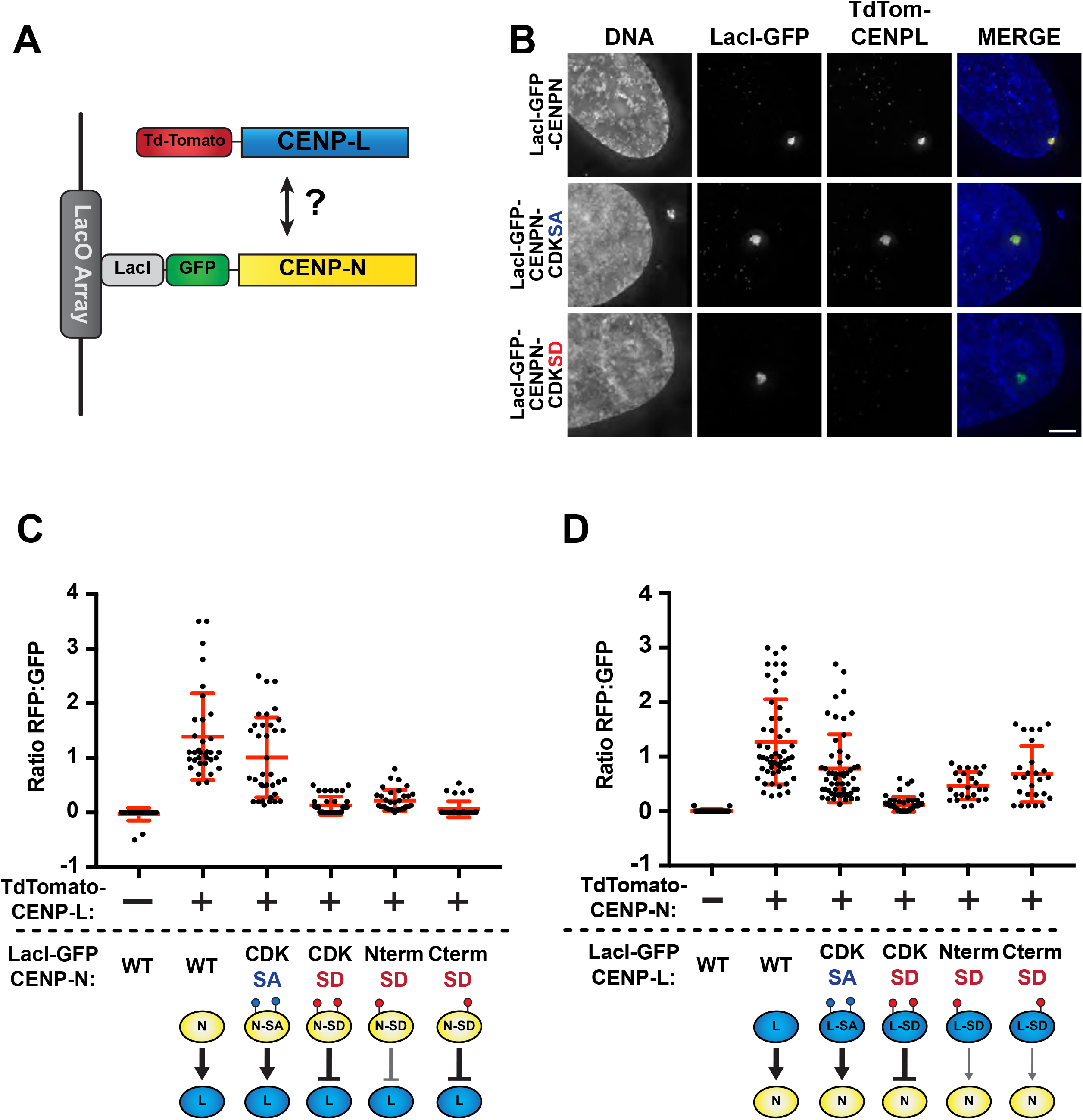
Phosphorylation state effects the interaction between CENP-N and CENP-L. **A.** Diagram illustrating the *lacO* ectopic targeting assay. **B.** Representative images of the Lac array assay for LacI-GFP-CENP-N and the recruitment of TdTomato-CENP-L. GFP and TdTomato images are scaled the same. Images were deconvolved. Scale bar is 5 μm. **C-D.** Quantification of fluorescence intensity for recruitment of TdTomato-tagged constructs to the *lacO* array by the LacI-GFP bait protein. Each dot represents the ratio between RFP:GFP fluorescence at a single foci. Error bars represent standard deviation.

### Phosphorylation of the CENP-N N-terminus disrupts its interaction with CENP-A

We next sought to evaluate whether phosphorylation affects additional CENP-L-N complex interactions. The association of CENP-N with kinetochores is dependent on its direct interaction with CENP-A (Carroll et al., 2009; McKinley et al., 2015). Structure-function analysis has defined the N-terminal region of CENP-N (aa 1-240) as both necessary and sufficient for its interaction with CENP-A and its localization to kinetochores during interphase (Carroll et al., 2009; Fang et al., 2015; McKinley et al., 2015). However, although the wild type N-terminal CENP-N fragment (CENP-N 1-240) localizes to kinetochores during interphase, a similar construct mimicking constitutive phosphorylation was unable to localize to kinetochores (Supplemental Fig. 2B). Similarly, a full length version of CENP-N with mutations in the phosphorylation sites within this N-terminal domain (T158D, T172D, T220D; referred to as CENP-N^NtermSD^; Fig. 4A) was unable to localize to kinetochores at any cell cycle stage (Fig. 4A) and was unable to rescue proper chromosome segregation in the absence of the endogenous protein (Fig. 4B-C). Together, this suggests that phosphorylation of the N-terminal CDK residues affects the interaction between CENP-N and CENP-A, preventing the recruitment of CENP-N to the centromere.

**Figure 4.**
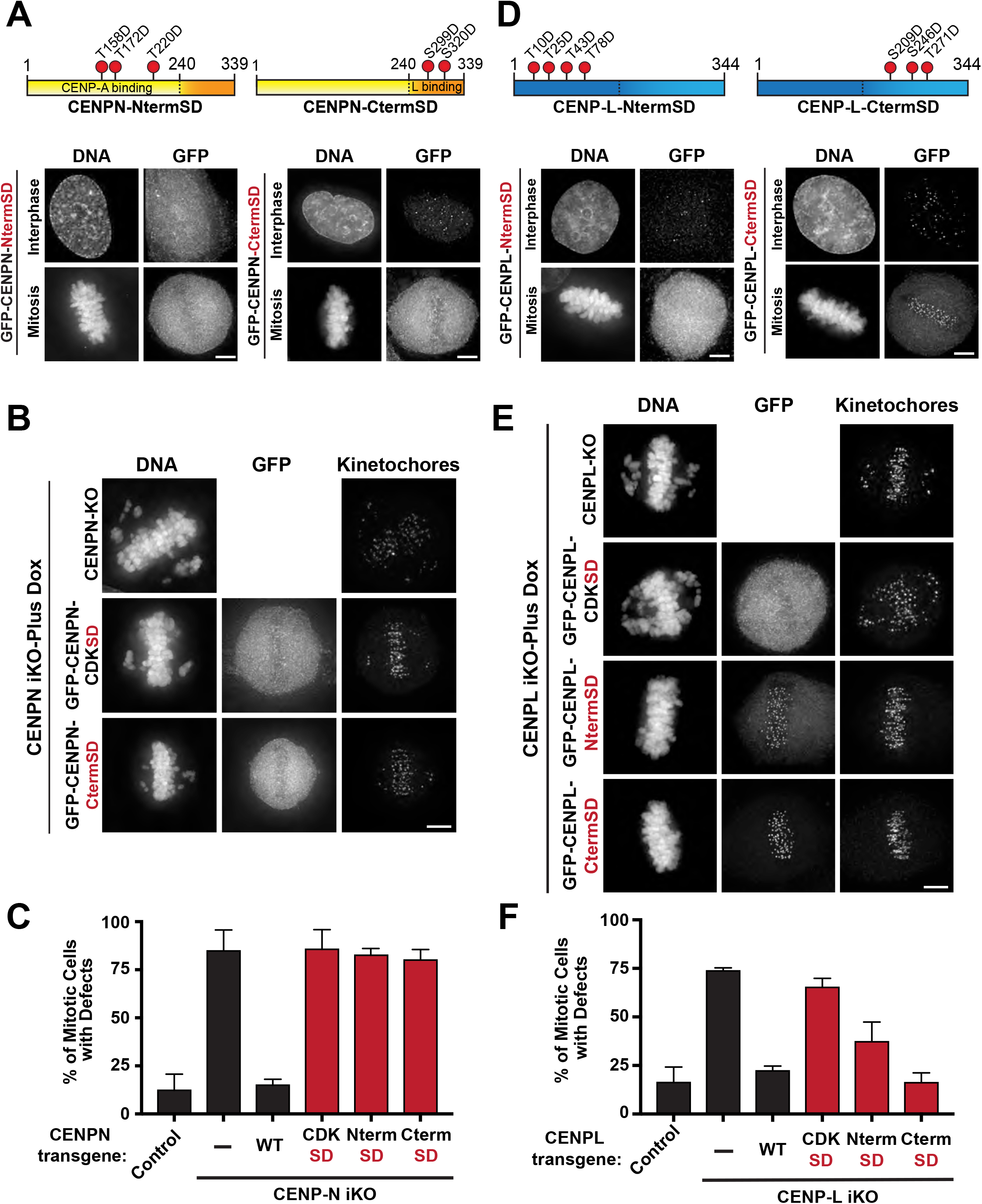
Splitting phosphomimetic point mutations between the N and C terminus results in differential localization behavior. **A.** Localization of CENP-N split phosphomutants during interphase and mitosis in the presence of endogenous protein. GFP-tagged constructs are transiently expressed for 48 hours before fixation. Diagrams illustrate what sites were mutated in each GFP-tagged construct. Scale bar is 5 μm. **B.** Representative images of mitotic phenotypes in replacement cells lines stably expressing the GFP tagged mutants after 4-day induction of CENP-N knockdown. Scale bar is 5 μM. **C.** Quantification of mitotic phenotypes in CENP-N split phosphomutant replacement cell lines. Approximately 100 cells were counted per condition. Note GFP-CENP-N-NtermSD replacement cell line is not a clonal cell line, and it only enriched for GFP fluorescence. All other GFP rescue cell lines are clonal. **D.** Localization of CENP-L split phosphomutants during interphase and mitosis in the presence of endogenous protein. GFP-tagged constructs are transiently expressed for 48 hours before fixation. Diagrams illustrate what sites were mutated in each mutant construct. Scale bar is 5 μm. **E.** Representative images of mitotic phenotypes in replacement cells lines stably expressing the GFP tagged mutants after 4-day induction of CENP-N knockdown. All GFP replacement cell lines are clonal. Scale bar is 5 μm. **F.** Quantification of mitotic phenotypes in CENP-L split phosphomutant replacement cell lines. Approximately 100 cells were counted per condition.

### Phosphorylation of the CENP-N C-terminus regulate CENP-N-CENP-L interactions

The C-terminus of CENP-N is necessary to mediate its interaction with CENP-L and is required to facilitate CENP-N kinetochore localization during mitosis (Carroll et al., 2009; Hinshaw and Harrison, 2013; McKinley et al., 2015; Pentakota et al., 2017). To test how phosphorylation affects the CENP-N-CENP-L interaction, we generated CENP-N mutants that specifically target its C terminal region: CENP-N^CtermSD^: S299D, S320D (Fig. 4A). The CENP-N^CtermSD^ mutant displayed differential cell cycle-dependent localization to kinetochores. Although the CENP-N^CtermSD^ mutant maintained its ability to localize to kinetochores during interphase, this mutant did not localize to kinetochores in mitosis. This altered temporal localization recapitulates the localization behavior of truncating the CENP-N C-terminus (Δ240-339), supporting the functional requirement of the C-terminal region of CENP-N in maintaining its kinetochore localization in mitosis (Supplemental Fig. 2C). Consistent with this localization behavior, this mutant failed to restore mitotic function in the absence of endogenous CENP-N (Fig. 4B-C) and failed to associate with CENP-L in the *lacO* tethering assay (Fig. 3C). Interestingly, we found that the CENP-N^NtermSD^ mutant also interacted poorly with CENP-L (Fig. 3C) suggesting that this mutant is defective in its binding to both CENP-A and CENP-L. Together, these observations highlight the requirement for the interaction between CENP-L and the C-terminal region of CENP-N to mediate kinetochore localization in mitosis, but not interphase.

### Phosphorylation sites in both the CENP-L N-terminal and C-terminal regions regulate its interaction with CENP-N

CENP-L interacts with both CENP-N and other CCAN components, but its functional domains are less clearly defined. Previous work utilizing structural predictions for CENP-L, as well as comparisons to the structure of the yeast ortholog, have suggested that the N-terminus of CENP-L has structural homology to CENP-N whereas the C-terminus mediates its interaction with CENP-N (Hinshaw and Harrison, 2013; Pentakota et al., 2017). To test the functional contributions of each domain of CENP-L, we generated phospho-mimetic mutants targeting the CDK sites in either the N or C terminal regions of CENP-L. Given the distribution of the phosphorylation sites within the protein and secondary structure predictions, we defined the N-terminal domain of CENP-L as amino acids 1-172 (CENP-L^NtermSD^) and the C-terminal region as amino acids 173-344 (CENP-L^CtermSD^) (Fig. 4D). Interestingly, the two split CDKSD mutants displayed distinct localization behaviors. In the presence of endogenous CENP-L, the CENP-L^NtermSD^ mutant displayed reduced localization to kinetochores during interphase and failed to localize in mitosis (Fig. 4D). Consistent with this altered localization, the CENP-L^NtermSD^ mutant showed reduced recruitment of CENP-N in the *lacO*/LacI assay, although this mutant was less defective than the full CENP-L^CDKSD^ mutant (Fig. 3D). However, kinetochore localization for the CENP-L^NtermSD^ mutant was restored in both interphase and mitosis following depletion of the endogenous protein (Fig. 4E) and was able to largely rescue chromosome mis-segregation (Fig. 4F). In contrast, the CENP-L^CtermSD^ mutant was not defective in its ability to localize or rescue mitotic phenotypes in the replacement cell line (Fig. 4D-F). Additionally, the CENP-L^CtermSD^ mutant recruited CENP-N to the *lacO* array in the *lacO*/LacI-binding assay similar to wild type CENP-L (Fig. 3D). Given that both CENP-L split mutants are able to interact with CENP-N to different degrees, this data suggests that both the N-terminus and C-terminus of CENP-L contribute to its interaction with CENP-N, but that the CENP-L N-terminus is particularly important for its localization in mitosis.

### Disrupting phosphorylation prevents CENP-LN complex kinetochore targeting

The results described above suggest that mimicking constitutive phosphorylation of either CENP-L or CENP-N is sufficient to disrupt CENP-LN complex interactions and kinetochore localization. In contrast, preventing the phosphorylation of either individual protein did not result in significant defects in CENP-LN complex formation or function. One possible explanation for this discrepancy is that phosphorylation of one subunit of this complex is sufficient to regulate the CENP-L-CENP-N interaction. Indeed, phospho-mimetic CENP-L^CDKSD^ mutants are unable to interact with either wild type CENP-N or the CENP-N^CDKSA^ mutants in the *lacO* ectopic targeting assay (Supplemental Fig. 3A). Therefore, to assess the consequences of failing to regulate the CENP-LN complex downstream of CDK, we hypothesized that it would be necessary to simultaneously prevent the phosphorylation of both CENP-N and CENP-L. Despite repeated attempts, we were unable to isolate cell lines stably co-expressing CENP-N^CDKSA^ and CENP-L^CDKSA^ (data not shown). Thus, to circumvent potential technical challenges associated with expressing both mutants constitutively, we developed a conditional strategy. For these experiments, we stably expressed either CENP-N^CDKSA^ or CENP-L^CDKSA^ individually in a double inducible knockout cell line expressing guides targeting both CENP-N and CENP-L. Following 3 days of Cas9 induction to deplete the endogenous protein, we then transiently expressed the reciprocal binding partner and assessed kinetochore localization and mitotic phenotypes 48 hours later.

In cell lines stably expressing either GFP-CENP-N or the GFP-CENP-N^CDKSA^ mutant, transient transfection of wild-type TdTomato-CENP-L restored localization of both GFP-CENP-N and GFP-CENP-N^CDKSA^ (Fig. 5A-B). Similarly, transiently expressing the TdTomato-CENPL^CDKSA^ mutant restored localization and rescued mitotic function in cell lines stably expressing the wild type GFP-CENP-N. In contrast, combining the TdTomato-CENPL^CDKSA^ and GFP-CENP-N^CDKSA^ mutants resulted in the failure of either protein to localize to kinetochores and severe mitotic defects (Fig. 5A-B). We observed similar results in experiments using cell lines stably expressing either CENP-L or the CENP-L^CDKSA^ mutant with the transient introduction of CENP-N constructs (Fig. 5A). Importantly, although combining the CENP-N^CDKSA^ and CENP-L^CDKSA^ mutants in the replacement assay prevented kinetochore localization, these mutants were able to interact with each other in our *lacO* ectopic targeting assay, although to a lesser extent than the wild type proteins (Supplemental Fig. 3A). This suggests that the absence of proper kinetochore localization is due to a defect in the loading or recruitment of the CENP-N^CDKSA^/CENP-L^CDKSA^ complex to kinetochores. Finally, to dissect the minimal requirements for this synergistic defect in the absence of CDK phosphorylation, we tested split CDK-SA mutants, targeting a subset of phosphorylation sites in each protein (Fig. 4A, 4D). In our dual replacement assay, CENP-N split mutants were able to largely restore the localization of the stably expressed CENP-L^CDKSA^ mutant and rescue mitotic function (Fig. 5D), suggesting that the phosphorylation of both regions is important for the observed effects. We found that preventing phosphorylation of the CENP-L N-terminus (CENP-L^Nterm-CDKSA^) failed to rescue localization and mitotic function in cell lines stably expressing CENP-N-CDKSA (Fig. 5C). In contrast, the CENP-L^Cterm-CDKSA^ mutant fully rescued mitotic function and localization of the CENP-N^CDKSA^ mutant (Fig. 5C). This suggests that the N-terminus of CENP-L plays a critical role in regulating its interaction with CENP-N, and that the regulation of this interaction is important for CENP-LN complex localization. Thus, regulating the interaction between CENP-L and CENP-N is critical to ensure proper CENP-LN localization to kinetochores throughout the cell cycle.

**Figure 5.**
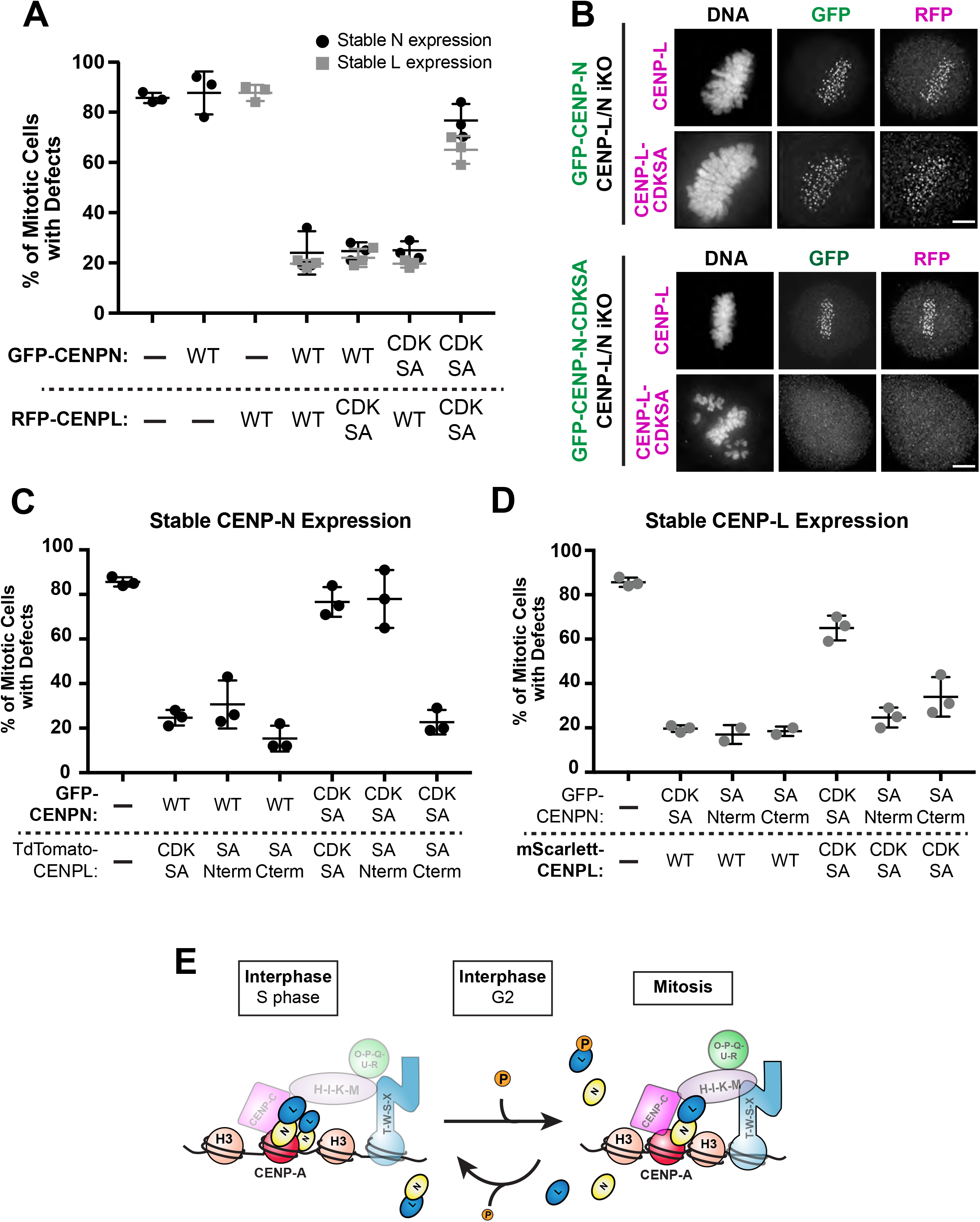
Preventing phosphorylation of both proteins negatively affects the function of the CENP-LN complex. **A.** Quantification of mitotic phenotypes in dual replacement cell lines following 4 days of Cas9 mediated knockdown of endogenous CENP-L and CENP-N and transfection of binding partner. Approximately 75 cells were counted per condition. **B.** Representative immunofluorescence images of cells in each condition. Cell lines are stably expressing GFP-CENP-N or GFP-CENP-N-CDKSA, both were transfected with either TdTomato-CENP-L or TdTomato-CENP-L-CDKSA. Scale bar is 5 μm. **C.** Quantification of mitotic phenotypes in dual replacement cell lines following 4 days total of Cas9 induction and knockdown of endogenous CENP-N. GFP-CENP-N or GFP-CENPN-CDKSA are stably expressed in this experiment. Tdtomato-CENP-L and TdTomato-CENP-L-CDKSA constructs were then transfected into dual replacement cell lines following 48 hours of dox induction. Media was then changed 16 hours post-transfection. Cells were fixed and processed 48 hours post transfection. Approx. 75 cells were quantified per condition for each experiment. **D.** Quantification of mitotic phenotypes in dual replacement cell lines following 4 days total of Cas9 induction and knockdown of endogenous CENP-L. mScarlett-CENP-L or mScarlett-CENPL-CDKSA are stably expressed in this experiment. GFP-CENP-N or GFP-CENP-N-CDKSA constructs were then transfected into dual replacement cell lines following 48 hours of dox induction. Media was then changed 16 hours post-transfection. Cells were fixed and processed 48 hours post transfection. Approx. 75 cells were quantified per condition for each experiment. **E.** Model Figure. Phosphorylation functions to weaken the interaction between CENP-L and CENP-N. Based on our data, the CENP-LN complex is phosphorylated by CDK upon entry into mitosis. This phosphorylation event helps to break the interaction between CENP-N and CENP-L. Following phosphorylation in mitosis, CENP-L and CENP-N must then be dephosphorylated to localize to the centromere.

### Cycles of dynamic phosphorylation drive inner kinetochore reorganization

Together, our data support a model in which dynamic cycles of phosphorylation act to regulate the recruitment of CENP-N and CENP-L to kinetochores in a cell cycle-dependent manner. Current structural models propose that there are sub-stoichiometric amounts of CENP-N during mitosis (Allu et al., 2019), but the basis for this change was unclear. To explain this behavior, prior studies suggested that the association of CENP-N with kinetochores is determined by the CENP-A chromatin state (Allu et al., 2019; Carroll et al., 2009; Fang et al., 2015). Indeed, CENP-N has been shown to have lower affinity for compact CENP-A chromatin *in vitro* (Fang et al., 2015). However, this model does not consider the functional importance of the interaction between CENP-N and CENP-L at kinetochores throughout the cell cycle, which mediates interactions with other CCAN components. Our work demonstrates that CENP-N-CENP-L interactions, which are necessary for CENP-N localization in mitosis, are regulated in a cell cycle-dependent manner. We find that phosphorylation of either CENP-L or CENP-N disrupts the interaction between these two proteins and thus the localization of this complex to kinetochores. Therefore, phosphorylation of a subset of the CENP-L/N complex molecules would result in a decrease in the total amount of CENP-L/N molecules at the kinetochore as cells enter mitosis, essentially acting as a molecular “pruning” event. Importantly, dynamic cycles of phosphorylation and dephosphorylation are critical for proper CENP-LN localization, as simultaneously preventing phosphorylation of CENP-L and CENP-N also negatively affects the behavior of this complex in the cell. We propose that the recruitment of this complex to kinetochores during S phase requires this mitotic phosphorylation followed by the subsequent dephosphorylation of both CENP-L and CENP-N to enable their interaction and loading at centromeres (Fig. 5E). Thus, despite the constitutive presence of the CENP-LN complex at centromeres, this dynamic regulatory control allows the inner kinetochore to set different organization and assembly states throughout the cell cycle.

## Acknowledgements

The authors thank the members of the Cheeseman lab for their suggestions and contributions to this project. This work was supported by grants from NIH/National Institute of General Medical Sciences (R35GM126930) and the NSF (2029868), to IMC and a National Science Foundation Graduate Research Fellowship to A.P.N.

## Methods

### Molecular Biology

The CENP-L targeting sgRNA (TCTGCAGAAACGATTAGAAT) was cloned into pLenti-sgRNA with either puro resistance or blast resistance. All GFP-tagged constructs used for expression in replacement cell lines were generated by cloning codon optimized guide hardened human cDNA into a pBABE-blast vector (pIC242) containing an N-terminal LAP tag (GFP-TEV-S) as described in (Cheeseman and Desai, 2005). The CENP-N and CENP-L split mutants were constructed by two rounds of overlap PCR using both the wild type and mutant cDNA’s. Dual replacement CENP-N and CENP-L expressing transgene was generated by synthesizing a mEGFP-MCS-T2A-mScarlett-MCS gene fragment and cloning this into a pLenti vector with a Hygromycin resistance cassette. cDNA’s for codon optimized cDNA’s were then cloned into this plasmid described previously.

### Cell Culture Conditions

HeLa cell lines were cultured in Dulbecco’s modified Eagle medium (DMEM) supplemented with 10% fetal bovine serum (FBS), 100 U/mL penicillin/streptomycin, and 2 mM L-glutamine at 37°C with 5% CO_2_. Inducible knockout cell lines were similarly cultured except the media was supplemented with certified tetracycline free FBS. Transient transfections were performed with Xtremegene-9 and Optimem as per manufacturer’s instructions. Transfected cells were fixed and processed 48 hours post transfection. The U2OS *lacO* array cell lines were cultured in similar media to Hela and maintained in 200 ug/mL Hygromycin.

### Replacement Cell Line Generation

Replacement cell lines were generated using the inducible CRISPR Cas9 HeLa cell line generated previously as described in (McKinley et al., 2015). The CENP-N knockout cell line was described previously in (McKinley et al., 2015). The CENP-L knock-out cell line was generated by introducing the CENP-L targeting guide into the inducible Cas9 cell line via lentiviral transduction as described in (McKinley et al., 2015). Single replacement cell lines were generated by introducing GFP-tagged constructs into their corresponding knockout cell lines by retroviral transfection followed by selection with 2ug/mL Blasticidin. Cells were then sorted for single clones. Where noted, cell lines enriched for fluorescence were not single cell sorted and instead collected based on positive fluorescent signal. The CENP-LN dual knockout cell line was generated by introducing the CENP-L guide cloned into a pLenti guide plasmid with blast resistance into the CENP-N ko cell line via lentiviral transduction. Cells were then selected with both 0.5 μg/mL puromycin and 2 μg/mL blasticidin for 2 weeks. The dual replacement transgene was also introduced via lentiviral transduction, cells were then selected with 0.5 μg/mL puromycin, 2 μg/mL blasticidin, and 400 μg/mL hygromycin. Cells were then sorted for both single cell clones and a fluorescently enriched population of cells.

To induce Cas9 expression in the inducible knock-out cell lines, cells were treated with 1ug/mL doxycycline at 0h, 24h, and 48h. Cells were then fixed and processed at 96h (on day 4). For transfection into dual replacement cell lines, cells were transfected on the third day of dox induced Cas9 expression using Xtremegene (at the same time as addition of dox at 48 hrs post induction). Media was changed 16 hours post transfection. Cells were fixed 48 hours post transfection.

### Immunofluorescence and Microscopy

Cells for immunofluorescence were seeded on glass coverslips coated with poly-L-lysine (Sigma-Aldrich). For all experiments, except those determining kinetochore levels throughout the cell cycle, cells were fixed in 4% formaldehyde (Sigma-Aldrich) diluted in PBS for 10 minutes. Coverslips were washed with PBS plus 0.1% Triton X-100 (PBS-TX). Blocking and primary antibody dilutions were performed in Abdil (20 mM Tris, 150 mM NaCl, 0.1% Triton X-100, 3% BSA and 0.1% NaN3, pH 7.5). GFP-booster (Chromotek; 1:300 dilution) was used to amplify GFP fluorescence for all tagged transgenes. Kinetochores were detected using anticentromere antibody (ACA) (Antibodies Inc #15234; 1:100 dilution). For assessment of mitotic phenotypes, microtubules were stained using DM1α (Sigma-Aldrich; 1:3000 dilution). For assessment of protein localization in the dual replacement cell lines, the RFP signal was amplified with an anti-RFP antibody (Rockland; 1:3000 dilution). Cy2-, Cy3-, and Cy5-conjugated secondary antibodies (Jackson ImmunoResearch Laboratories) were used at a 1:200 dilution in PBS plus 0.1% Triton X-100. DNA was visualized by incubating cells in 1 ug/mL Hoechst33342 (Sigma-Aldrich) in PBS plus 0.1% Triton X-100 for 10 min. Coverslips were mounted using PPDM (0.5% p-phenylenediamine and 20 mM Tris-Cl, pH 8.8, in 90% glycerol) and sealed with nail polish. For cell cycle quantifications the following antibodies were used: anti-CENPC (Gascoigne et al., 2011); anti-PCNA (Abcam; 1:500 dilution); anti-Cyclin B (Santa Cruz; 1:500 dilution). For GFP-tagged quantifications cells were fixed in 4% formaldehyde diluted in PBS and then treated with MeOH for 2 min at −20C. For CENP-C quantification, cells were fixed in 4% formaldehyde diluted in PBSTx and then treated with MeOH for 2 min at −20C. Images were acquired on a DeltaVision Core deconvolution microscope (Applied Precision) equipped with a CoolSnap HQ2 charge-coupled device camera (Photometrics). All images were maximally projected and deconvolved.

For quantification of fluorescence intensity, maximum intensity projections were generated using the Deltavision Softworx Software (GE Healthcare). Integrated fluorescence intensity was measured using the MetaMorph software (Molecular Devices). For fluorescence intensity measurements at kinetochores, a 7×7 pixel region surrounding a kinetochore and a region in the cytoplasm were selected. Background measurements were subtracted from each kinetochore measurement. 5 cells were imaged per cell cycle stage and approx. 40 kinetochores were imaged per cell. All values measured for kinetochores at G2 were divided by 2 because individual kinetochore cannot be resolved. For Lac array experiments, a 10×10 pixel regions surrounding a foci and a region of the cytoplasm were selected. Background measurements were subtracted from each foci measurement. Approx. 10 cells were quantified per condition.

### Immunoprecipitation and Mass Spectrometry

For mitotically enriched immunoprecipitations, cells from 40x 15cm plates were arrested with 330nM nocodazole (Sigma-Aldrich) for 16 hours and harvested by shake-off. For cells enriched in S-phase, cells from 40x 15 cm plates were treated with 2mM thymidine (Sigma-Aldrich) for 18 hours and harvested with PBS plus 5mM EDTA. Immunoprecipitations for mass spectrometry analysis were performed as described previously ((Cheeseman and Desai, 2005). Harvested cells were washed in PBS and resuspended 1:1 in 1X Lysis Buffer (50 mM HEPES, 1 mM EGTA, 1 mM MgCl2, 100 mM KCL, 10% glycerol, pH 7.4) then drop frozen in liquid nitrogen. Cells were thawed after addition of an equal volume of 1.5X lysis buffer supplemented with 0.075% Nonidet P-40, 1X Complete EDTA-free protease inhibitor cocktail (Roche), 1 mM PMSF, 20 mM beta-glycerophosphate, 1 mM sodium fluoride, and 0.4 mM sodium orthovanadate. Cells were lysed by sonication and cleared by centrifugation. The supernatant was mixed with Protein A beads coupled to rabbit anti-GFP antibody (Cheeseman and Desai, 2005) and placed on a rotating wheel at 4°C for 1hr. Next, the beads were washed five times in 50 mM HEPES, 1 mM EGTA, 1 mM MgCl2, 300 mM KCl, 10% glycerol, 0.05% NP-40, 1 mM DTT, 10 μg/mL leupeptin/pepstatin/chymostatin, pH 7.4 wash buffer (wash buffer). Following a final wash in wash buffer without detergent, bound protein was eluted with 100 mM glycine pH2.6. Eluted proteins were precipitated by addition of 1/5^th^ volume trichloroacetic acid at 4°C overnight. Precipitated proteins were reduced with TCEP, alkylated with iodoacetamide, and digested with mass-spectrometry grade Lys-C and trypsin (Promega). Digested peptides were purified using C18 spin columns (Pierce) according to the manufacturer’s instructions. Samples were analyzed on an LTQ XL Ion Trap mass spectrometer (Thermo Fisher) coupled with a reverse phase gradient over C18 resin. Data were analyzed using Proteome Discoverer (Thermo Fisher).

## Supplemental Figures

**Supplemental Figure 1.**
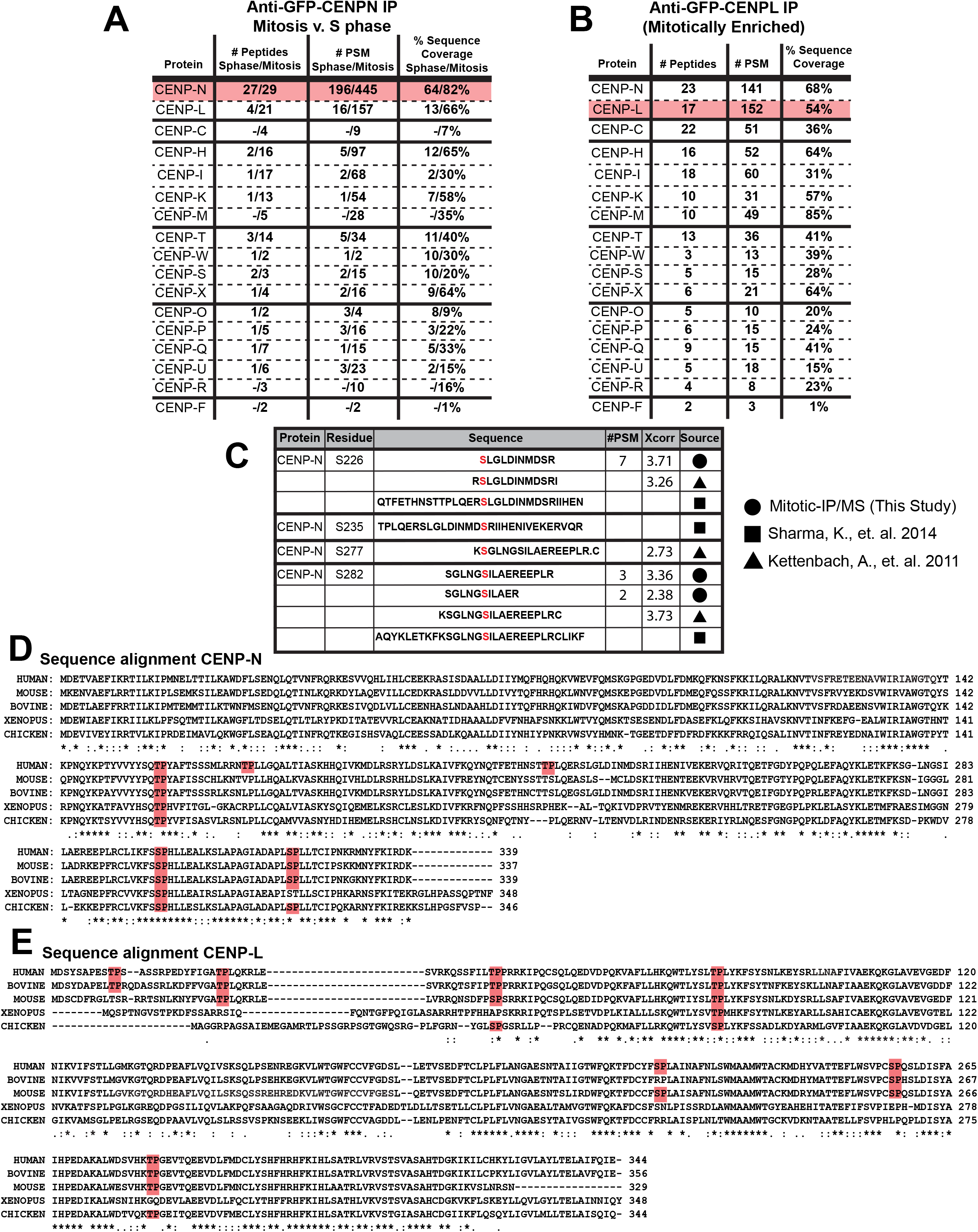
Summary of mass spec data and conservation of CDK phosphorylation sites in CENP-L and CENP-N. **A-B.** Summary of proteins identified by mass spectrometry after each indicated immunoprecipitation. The proteins listed are all other inner kinetochore components with the corresponding peptide count, #PSMs, and % Sequence coverage. The bait for each IP is highlighted in red. In CENP-N chart values for both S phase enrichment and Mitosis enrichment are shown. **C.** Table representing the non-CDK sites identified by our mass spec analysis and previously reported in the literature. **D.** Sequence alignments for the SP/TP sites found throughout the CENP-N sequence. SP/TP sites are highlighted in red. Sequence alignments generated with UniProt alignment. **E.** Sequence alignments for the SP/TP sites found throughout the CENP-L sequence. SP/TP sites are highlighted in red. Sequence alignments generated with UniProt alignment.

**Supplemental Figure 2.**
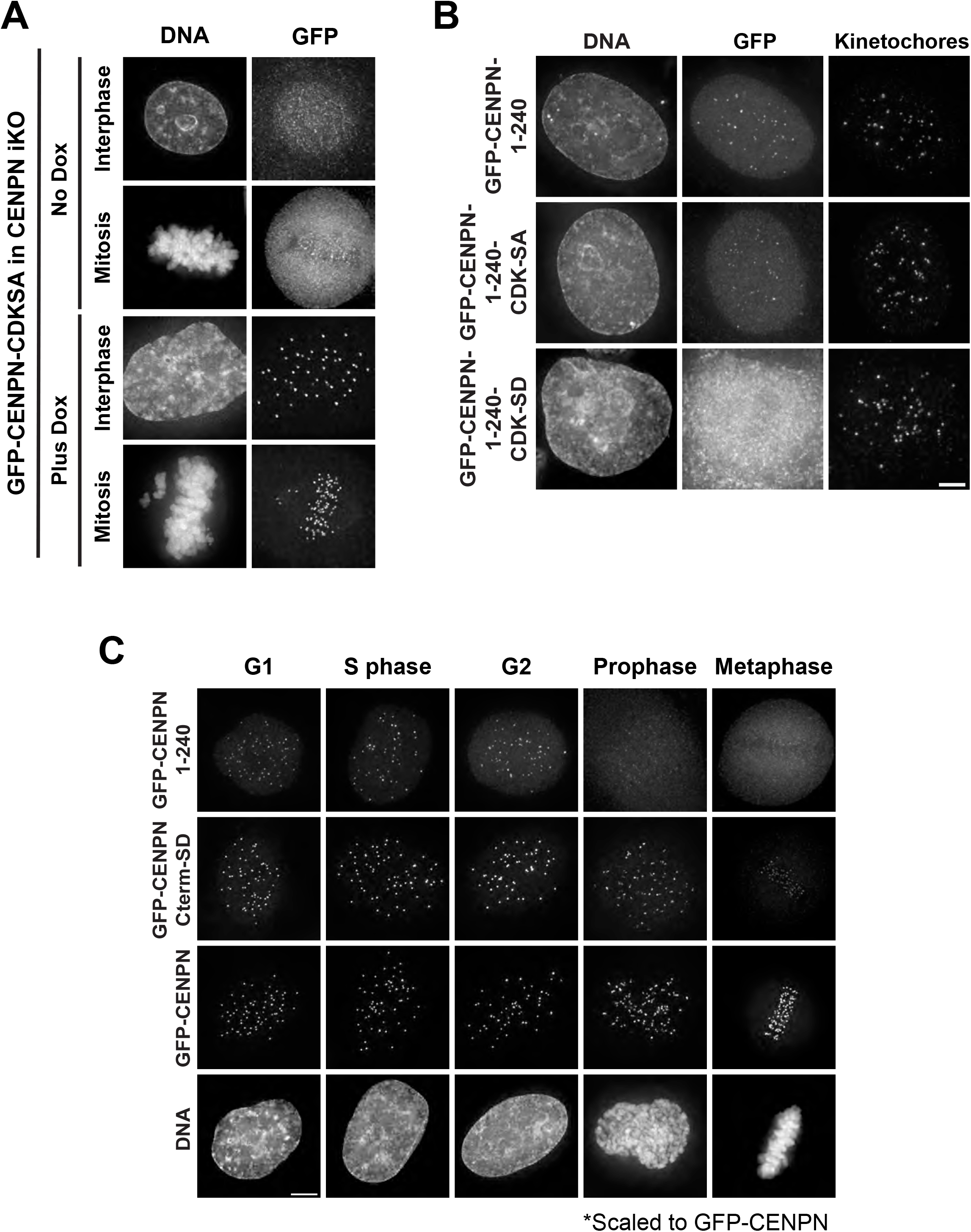
Phosphorylation state affects the localization of CENP-N. **A.** Representative images of GFP-CENP-N-CDKSA localization in the CENP-N iKO cell lines. Top panel shows localization in the absence of Cas9 expression. The bottom panel shows localization following 4 day induction of Cas9 expression and therefore depletion of endogenous protein. Scale bar 5 μm. **B.** Images showing the kinetochore localization of the GFP-CENP-N-1-240 phosphomutants. Constructs were transiently transfected into HeLa cells. Scale bar is 5 μm. **C.** Representative images of three GFP-CENP-N mutants and their kinetochore localization throughout the cell cycle from cell lines stably expressing these constructs. All images are scaled relative to GFP-CENP-N signal. DNA images shown are from the GFP-CENP-N cells. Scale bar is 5μm.

**Supplemental Figure 3.**
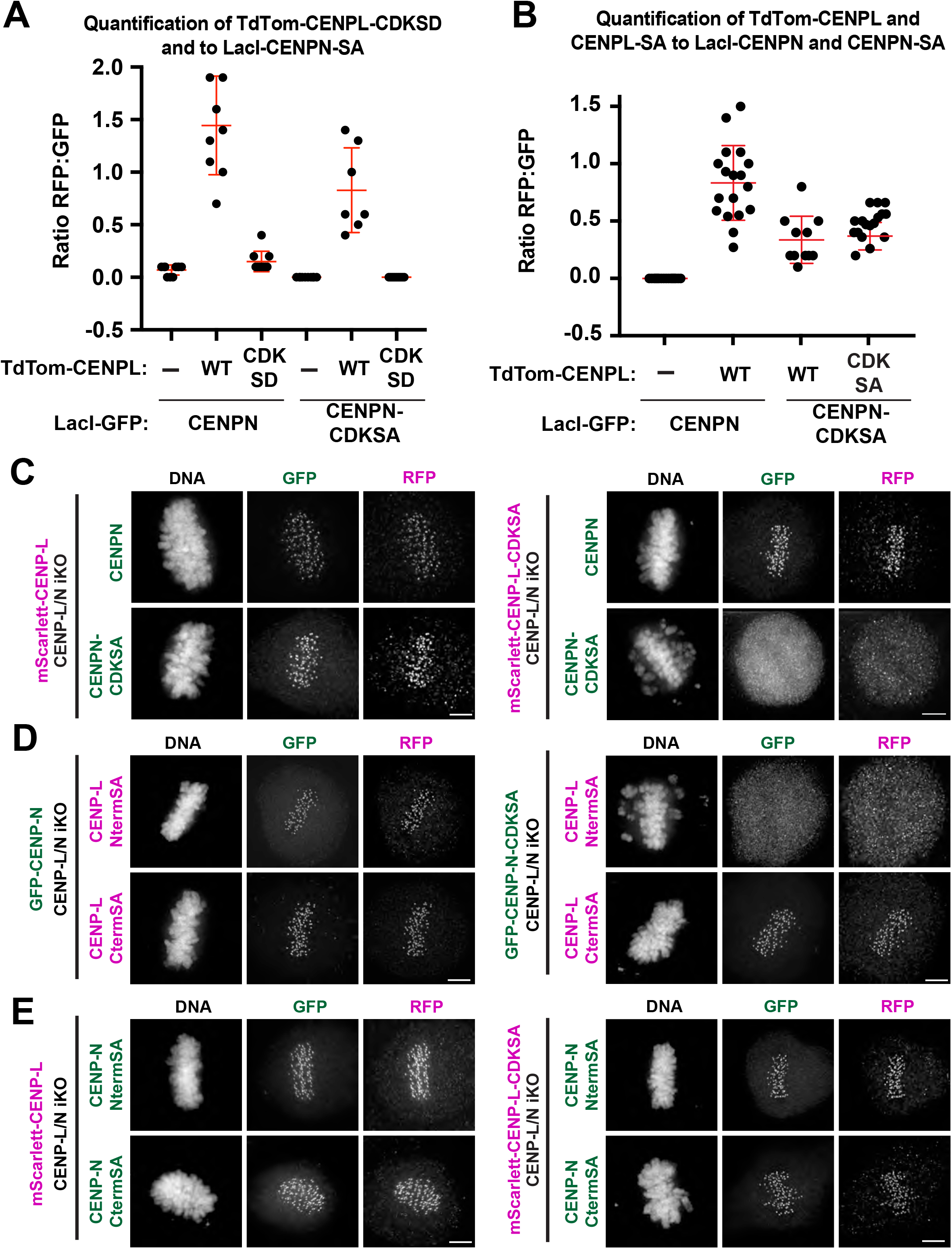
Preventing phosphorylation of CENP-L and CENP-N affects the recruitment of this complex to the kinetochore but not their ability to interact. **A.** Quantification of fluorescence intensities for recruitment of TdTomato-tagged CENP-L or CENP-L-CDKSD to the LacI-CENP-N/NSA foci. Each dot represents the ratio between TdTomato:GFP fluorescence at a single foci. Data combined from three individual experiments. Error bars represent standard deviation. **B.** Quantification of fluorescence intensities for recruitment of TdTomato-tagged CENP-L or CENP-LSA to the LacI-CENP-N/NSA foci. Each dot represents the ratio between TdTomato:GFP fluorescence at a single foci. Data combined from three individual experiments. Error bars represent standard deviation. **C.** Representative immunofluorescence images of cells in each condition. Cell lines are stably expressing mScarlett-CENP-L or mScarlett-CENP-L-CDKSA, both cell lines were transiently transfected with plasmids expressing either GFP-CENP-N or GFP-CENP-N-CDKSA. Cells were transfected 48 hours post dox induction of Cas9. Cells were fixed and processed for immunofluorescence 48 hours post transfection. Scale bar is 5 μM. **D.** Representative immunofluorescence images of cells in each condition. Cell lines are stably expressing GFP-CENP-N or GFP-CENP-N-CDKSA, both were transiently transfected with plasmids expressing TdTomato-CENP-L-Nterm-CDKSA or TdTomato-CENP-L-Cterm-CDKSA Cells were transfected 48 hours post dox induction of Cas9. Cells were fixed and processed for immunofluorescence 48 hours post transfection. Scale bar is 5 μm. **E.** Representative immunofluorescence images of cells in each condition. Cell lines are stably expressing mScarlett-CENP-L or mScarlett-CENP-L-CDKSA, both were transiently transfected with plasmids expressing GFP-CENP-N-Nterm-CDKSA or GFP-CENP-N-Cterm-CDKSA Cells were transfected 48 hours post dox induction of Cas9. Cells were then fixed and processed for immunofluorescence 48 hours post transfection. Scale bar is 5 μm.

